# Developmental dynamics of the prefrontal cortical SST and PV interneuron networks: Insights from the monkey highlight human-specific features

**DOI:** 10.1101/2024.07.10.602904

**Authors:** Nafiseh S. Hosseini Fin, Adrian Yip, Leon Teo, Jihane Homman-Ludiye, James A. Bourne

## Abstract

The primate prefrontal cortex (PFC) is a quintessential hub of cognitive functions. Amidst its intricate neural architecture, the interplay of distinct neuronal subtypes, notably parvalbumin (PV) and somatostatin (SST) interneurons (INs), emerge as a cornerstone in sculpting cortical circuitry and governing cognitive processes. While considerable strides have been made in elucidating the developmental trajectory of these neurons in rodent models, our understanding of their postmigration developmental dynamics in primates still needs to be studied. Disruptions to this developmental trajectory can compromise IN function, impairing signal gating and circuit modulation within cortical networks. This study examined the expression patterns of PV and SST, ion transporter KCC2, and ion channel subtypes Kv3.1b, and Nav1.1 -associated with morphophysiological stages of development in the postnatal marmoset monkey in different frontal cortical regions (granular areas 8aD, 8aV, 9, 46; agranular areas 11, 47L). Our results demonstrate that the maturation of PV+ INs extends into adolescence, characterized by discrete epochs associated with specific expression dynamics of ion channel subtypes. Interestingly, we observed a postnatal decrease in SST interneurons, contrasting with studies in rodents. This endeavor broadens our comprehension of primate cortical development and furnishes invaluable insights into the etiology and pathophysiology of neurodevelopmental disorders characterized by perturbations in PV and SST IN function.

**Summary Statement:** The prefrontal cortex (PFC) in primates is crucial for cognitive functions, with parvalbumin (PV) and somatostatin (SST) interneurons playing key roles. This study in marmoset monkeys explores their developmental dynamics, revealing prolonged maturation of PV interneurons and contrasting SST patterns from rodents, enhancing understanding of primate cortical development.

## Introduction

Distinguishing it from the rodent primate prefrontal cortex (PFC), specific areas of the primate, including the human PFC, possess a distinct granular layer [1, 2], resulting in the orchestration of particular functions and connectivity [3]. In addition to the expanded primate cortex, an evolved thalamic network, the medial pulvinar, is interconnected with these areas [4]. For example, the heteromodal dorsolateral prefrontal cortex (DLPFC) of the anthropoids comprises Broadman’s areas (BA) 8, 9, and 46, receiving graded input from multiple sensory and multimodal areas, with projections to and from occipital, temporal, and parietal lobes, and sensorimotor cortices. All the areas of the PFC are also interconnected through the expanded primate-specific medial pulvinar nucleus of the thalamus [5]. The evolution of the mosaic primate PFC has seen the number of discrete prefrontal areas increase in the anthropoids from 26 in marmosets (*Callithrix jacchus*) to 35 in macaques (*Macaca mulatta*), culminating with 45 in humans (*Homo sapiens)* [2].

The emergence of new brain areas, architectonics, and connectivity is not the only feature distinguishing the rodent neocortex from the primate. Primate evolution was realized by incorporating novel cell types in pre-existing brain regions [6]. The expansion of neuronal diversity in the primate brain disproportionately affects INs, suggesting they play a vital role in primate cognition. While INs account for 15–20 % of total cortical neurons in rodents, this ratio soars to 25–34% in primates [6, 7]. Therefore, understanding the establishment and physiology of primate-specific INs is a window into the evolution of primates’ unrivaled cognitive functions.

All the GABAergic INs found in adult primates, including humans, originate from the embryonic ganglionic eminences (GEs) in the subpallium, as well as the proliferative zones of the dorsal telencephalon, a primate-specific IN niche [8, 9]. The medial GE (MGE) is the source of two major classes of INs that express neuropeptide hormone, somatostatin (SST), or calcium-binding protein, parvalbumin (PV), referred to as SST+- and PV+ INs, respectively [10–12]. Putative SST+- and PV+ cells in the MGE are identified by the expression of the transcription factor SOX6 [9]. While it is not entirely elucidated in primates, rodent studies support that SST+ INs are generated early from asymmetrically dividing progenitors in the ventricular zone (VZ) [13, 14]. In contrast, PV+ INs are generated later from MGE subventricular zone (SVZ) progenitors that divide symmetrically [14]. The difference in birthdate and progenitor origin suggests these two subtypes perform distinct functions in developing and mature functioning neocortex. For example, mature PV+ INs tightly control pyramidal cell output. Thus, PV+ IN maturation in the PFC is crucial for cognitive development [15]. In contrast, rodent studies suggest a critical role of SST+ cells in establishing thalamic inputs during the early development of the barrel cortex [16]. This function remains to be investigated in primates.

As INs integrate their local network in the neocortex, they upregulate the expression of subtype markers such as calbindin (CB), PV, or SST. In addition, specific ion channels and transporters, including the potassium chloride cotransporter KCC2, sodium channel subtype Nav1.1, and potassium channel subtype Kv3.1b expression, are correlated with acquiring their mature functionality [17–22]. The upregulation of these proteins plays an essential role in IN homeostasis. PV and CB contribute to buffering the increasing calcium flux as the neurons become more active [23, 24]. The ion channels participate in the transition from an excitatory response to GABA to the conventional inhibitory response, as well as functional specialization of the IN-firing properties [25, 26]. The mature network is finally consolidated by depositing an elaborate scaffold of proteoglycans, collectively known as perineuronal nets (PNN) and myelination [27]. This marks the closure of maturation and, to some degree, plasticity and the inception of a functional adult network.

Maturation is a progressive phenomenon that occurs over a protracted period, especially in primates, from infancy for primary sensory areas to adolescence and adulthood for the PFC, as determined by longitudinal MRI analysis [28]. Utilizing the temporal expression of neurofilament proteins (nonphosphorylated neurofilament; NNF), restricted to a subset of mature excitatory neurons, we previously reported that the maturation of sensory networks was sequential and followed the processing hierarchy, with the primary visual cortex (V1) maturing first, followed by association cortices [29]. A similar approach to studying NNF in the PFC revealed a posterior-to-anterior maturation gradient, with the most anterior areas of the frontal pole developing the last [30].

Abnormal maturation or dysfunction of INs in the PFC has been linked to various neurodevelopmental disorders, including schizophrenia, autism spectrum disorders, and attention deficit hyperactivity disorder (ADHD) [31]. In particular, the evidence suggests that disrupting the delicate balance between excitatory and inhibitory neural activity in the PFC can lead to cognitive impairments and altered behavior [32]. Hence, understanding the molecular pathways involved in the maturation and functioning of INs is paramount in elucidating the underlying mechanisms that fail in neurodevelopmental disorders.

The marmoset monkey (*Callithrox jacchus*), with its phylogenetic proximity to humans and comparable cortical organization, offers a compelling avenue for investigating the nuanced development and maturation of PV and SST INs within the PFC [33]. Unraveling the ontogeny of these interneuronal populations in the marmoset, PFC holds profound implications for comprehending the evolutionarily conserved principles governing cortical circuit assembly and function, with potential translational relevance to human neurodevelopmental disorders [33].

This study seeks to illuminate the intricate choreography of PV and SST IN development and maturation within the marmoset PFC areas 8aD and V, 46, 9 (dorsolateral prefrontal cortex, granular cortex) and 11 (orbitofrontal cortex, dysgranular) and 47L (ventrolateral cortex, dysgranular). These areas were chosen as they span the anteroposterior and mediolateral axes of the PFC [34]. PV, SST, KCC2, Nav1.1, Kv3.1b, and PNN accumulation were profiled in each area from the early stages of postnatal life into adulthood. By integrating insights from molecular and cellular investigations, we aim to delineate the spatiotemporal dynamics, molecular determinants, and implications underlying the maturation of these key neuronal subsets, which may affect our understanding of neurodevelopmental disorders.

## Results

### Demarcation of areas 8aD, 8aV, 47L, 9, 46, and 11 in the marmoset PFC from infant to adult

The neocortex of the adult marmoset has been extensively mapped. Many resources are available to guide the demarcation of the adult neocortex. For example, [34]. However, for this study, we needed to extend this segmentation to the five postnatal developmental stages based on this study (Fig.1, A). The first observations were carried out on postnatal day 7 (PD7) when primary sensorimotor cortices express nonphosphorylated neurofilament (NNF), a marker of pyramidal neuron maturation [35]. Prefrontal areas show minimal NNF expression at that early stage, with only faint expression in areas 6D and 8aV in L5 at PD7 [30]. At one month postnatal (PM1), the expression of NNF remains faint in the prefrontal cortex. Still, it expands to secondary and tertiary sensory areas in both L3 and 5 [35] and has reached adult levels in motor areas [36]. At three months postnatal (PM3), our third stage of examination, maturation is complete in the motor cortex and advanced in the visual cortex. At the same time, the expression of NNF only now becomes apparent in the prefrontal areas [30]. The two subsequent stages explored, 9 and 12 months (PM9 and 12), corresponding to adolescence and the intensification of NNF expression in the prefrontal areas, including the DLPFC. The final stage, PM18, is adult, in which we expect all cortical areas to have reached peak maturation, according to the literature.

**Figure 1:**
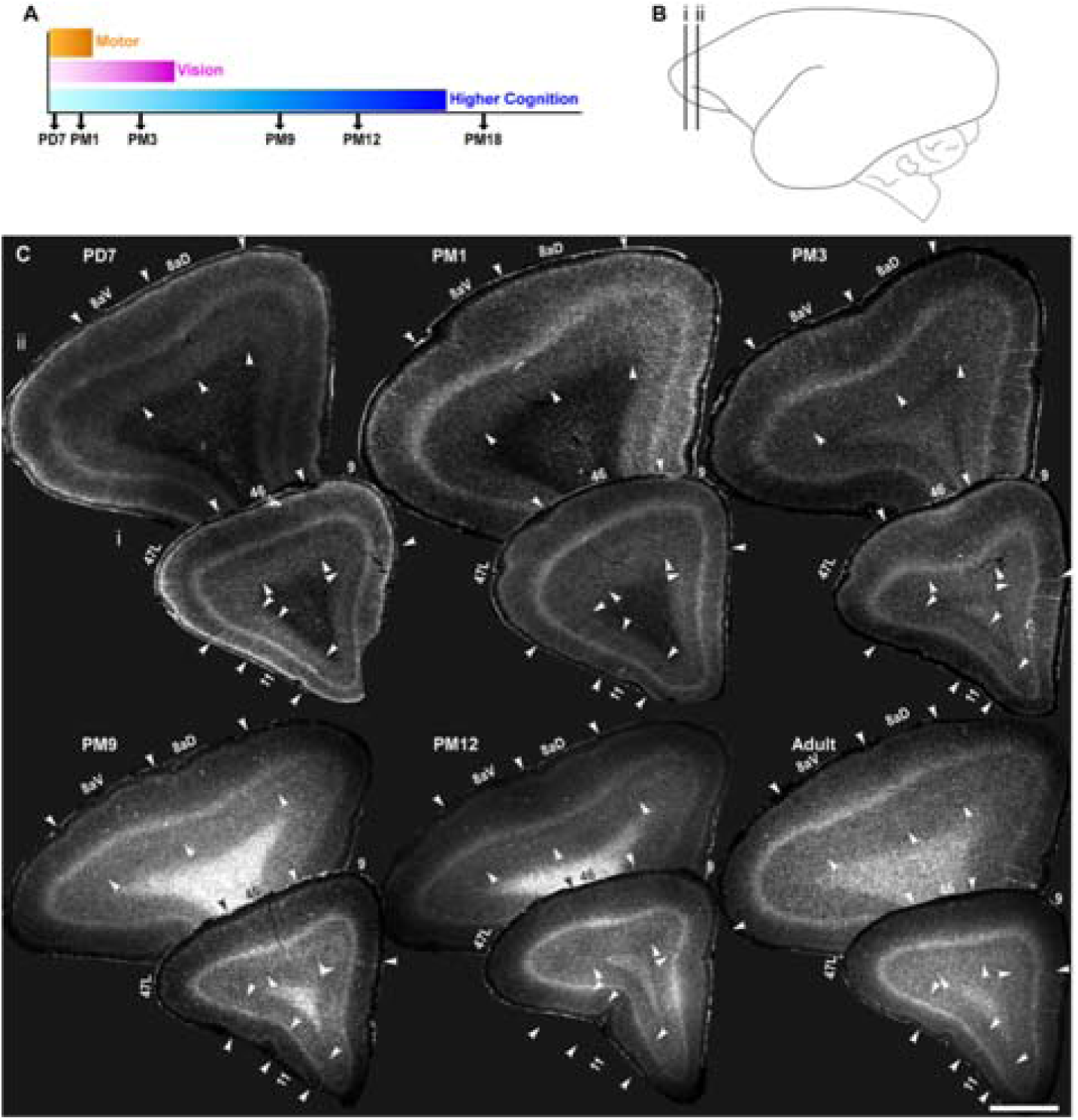
Cytoarchitecture of marmoset prefrontal neocortical areas 8aD, 8aV, 47L, 11, 9, and 46 from postnatal day (PD) 7 to adult. **A** Representation of developmental stages used in this study: postnatal day (PD) 7, postnatal month (PM) 1, 3,9, 12, and 18 presented in correlation with the indicative timing of the maturation of the cortical networks sustaining vision and higher cognitive and motor functions. **B** Position of the areas of interest represented on a schematic of the adult marmoset brain modified from the marmoset atlas [34]. **C** Representative examples of the laminar cytoarchitecture of the marmoset PFC coronal section stained with Hoechst from birth to adulthood. Changes in cortical layer thickness and density enable the demarcation of the borders between adjacent cortical areas, as indicated by the arrowheads. All the analyzed images were captured in the core region of the areas of interest for consistency within and across stages.

For this study, we focused on 6 PFC areas (Fig. 1, B). Areas 8, anterior dorsal (8aD) and anterior ventral (8aV), are part of the frontal eye fields (FEF) and participate in the control of saccadic eye movement [37]. While both 8aD and 8aV are components of DLPFC, they form a reference point in this study as their maturation has previously been explored using NNF as a proxy [30]. Moreover, compared to areas 9/ 46, their posterior position allows us to evaluate the hypothesis that cortical areas mature in a posterior-to-anterior sequence [38]. Finally, the analysis includes areas 47 (lateral prefrontal cortex; dysgranular) and 11 (orbitofrontal cortex; granular). These two areas fulfill different roles in cognitive flexibility and emotional and reward-based decision-making [39–42].

Due to its lissencephalic surface, the growth and postnatal expansion of the marmoset neocortex is relatively isotropic, allowing for the demarcation of individual cortices at various stages and comparison feasible. The precise cytoarchitectural boundaries of the areas of interest were demarcated based on the literature [30, 43, 44]. In short, variations in the evidence and thickness of L4 were the main point of reference for resolving the boundary between adjacent areas across all stages (Fig.1, C). For example, L4 in 8aV was characteristically thicker than in adjacent area 8aD, which could be delineated further by the relative expansion of L3 and 4, which had a sharp interface with L5 compared to neighboring areas. Area 46 possessed a well-defined but thinner L4 at the same mediolateral level. In contrast, area 47L, localized laterally to area 46, was characterized by a thinner L4 and thicker infragranular layers than area 46. Conversely, area 9 was identified to have a thin L4 and relatively homogeneous supra- and infragranular layers. Area 11, compared to the adjacent areas, was restricted by the thinnest L4 and occupied the anterior orbital sulcus.

### Synchronous postnatal decrease of SST+ interneurons in the prefrontal areas

Somatostatin (SST) is widely expressed across the brain, particularly in a subset of cortical INs. SST+ INs are one of the two subgroups of MGE-derived cortical INs, with the PV+ subtype comprising most cortical INs across mammals, including primates [6]. While the timing of incorporation of SST+ cells in neuronal networks in primates is not elucidated yet, rodent studies suggest an earlier integration of SST+ INs into functional networks, compared to PV+ cells [17, 45]. We observed SST+/ NeuN+ INs in the marmoset PFC as early as the first postnatal week in supragranular and infragranular layers, and the white matter (Fig. 2, A, B, and C), which had declined in all areas and layers by adolescence, (Fig. 2, D, and E). To substantiate our observation, we counted the density of SST+ in L3 and L5/6 for each time point. We normalized our values to the total number of neurons to account for cortical expansion due to the addition of glial cells and synapses during postnatal development. After confirming that the proteins were co-expressed within the same INs, we calculated the SST+/ NeuN+ neurons ratio in the field of view (Fig. 2, F-F”).

**Figure 2:**
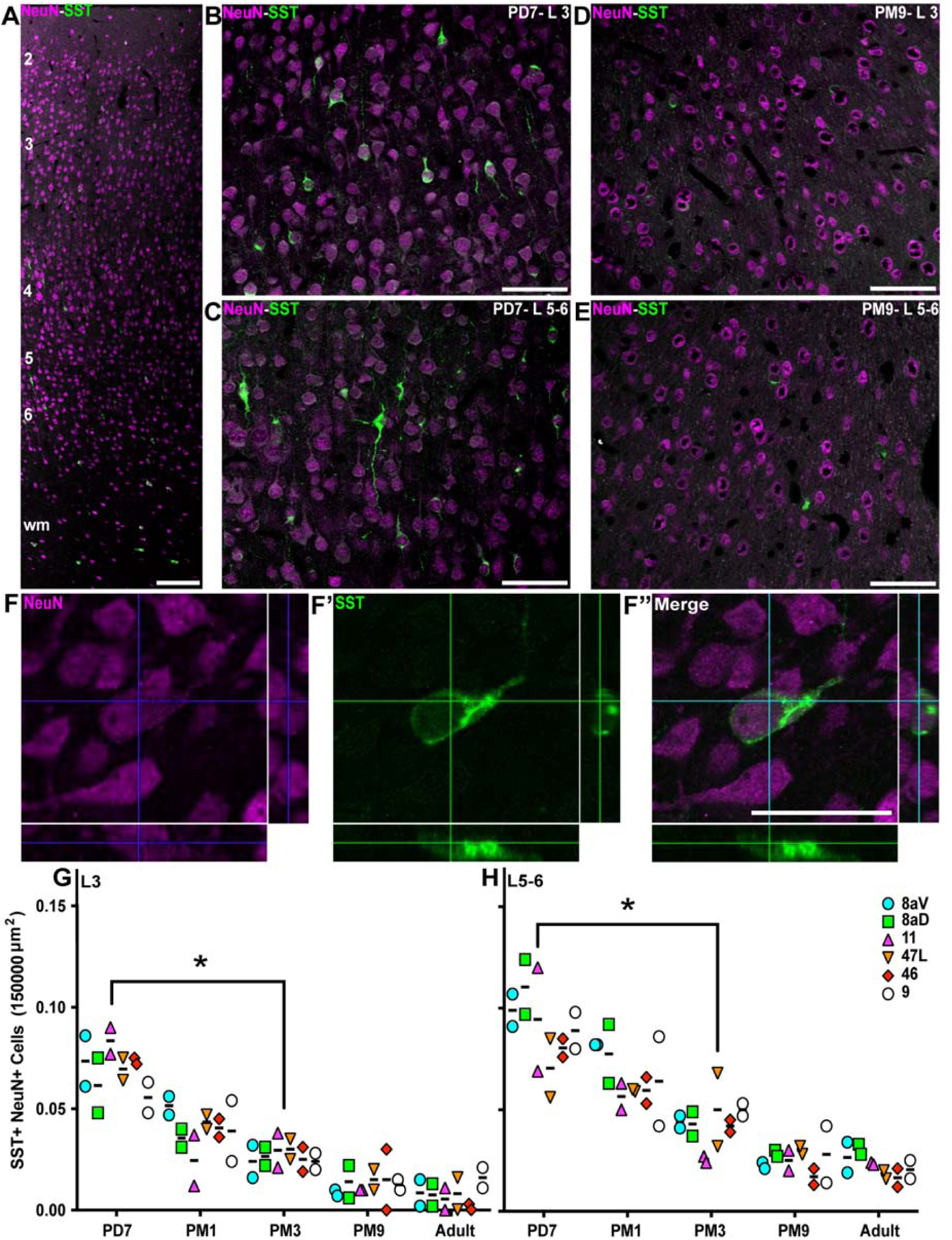
Progressive reduction of the proportion of SST+ interneurons in the marmoset PFC from birth to adolescence. SST+ interneurons (green) and NeuN+ neurons (magenta) are present across all cortical layers of the marmoset DLPFC (area 46) at PD7. Distribution of SST+ NeuN+ interneurons in area 46 L3 and 5/6 of at PD7 (**B**, **C**) and PM9 (**D**, **E**). **F-F”** Cutlines through a 3D stack acquired with a laser scanning confocal microscope confirm that NeuN nuclear labeling and cytoplasmic SST labeling correspond to a unique cell in area 46 L3 at PD7. The ratio of SST+ NeuN+ interneurons over the total number of NeuN+ neurons per 150,000 µm^2^ was calculated at each developmental stage in all 6 PFC areas of interest and plotted over time. The proportion of SST+ interneurons decreases progressively and at a comparable rate in all areas in the supragranular L3 (**G**) and L5/6 (**H**). Statistical analysis was performed using a nonparametric Kruskal–Wallis test followed by Dunn’s multiple comparisons. The data are presented as median ± interquartile ranges, * p<0.05: significant (n=2).

First, the relative distribution of SST across all PFC areas at each time point was insignificant. At PD7, SST+ INs accounted for 0.074 ± 0.02 of L3 neurons (Fig. 2, G). This value was similar across all areas of the prefrontal cortices examined. The ratio rapidly decreased over development, dropping to 0.027 ± 0.01 at PM3 (p≤0.01, Kruskal–Wallis test). The ratio of SST+ INs did not significantly vary during adolescence and adulthood, and the changes were homogenous across all PFC areas of interest. Analysis of L5/6 yielded a comparable profile (Fig. 2, H), with the exception that the ratio of SST+ over the total neuronal population was higher in the first postnatal week (0.09 ± 0.03) and decreased to (0.05+0.02, p≤0.01; Kruskal–Wallis test) at PM3.

To confirm the developmental reduction in SST+ INs did not result from apoptotic cell death at PM3 and more likely a down-regulation of the peptide, activated caspase 3 (aCasp3) labeling and pyknotic nuclear Hoechst stain was used. Results remained consistent between PD7 and PM3 in the supra- and infragranular layers of areas 46 and 47L (Sup. Fig. S1). Overall, these findings suggest that the maturation of SST+ INs is synchronized across the prefrontal cortex, unlike the sensory cortex. Considering this result, we investigated the developmental regulation of PV expression, the other primary class of cortical MGE-derived INs.

### A proportional increase in the fraction of PV+ interneurons in the prefrontal areas during preadolescence

PV+ INs were observed primarily in L3-5 (e.g., Fig. 3, A, area 46 at PM 9). At PD7, PV+ INs were largely absent from L3 (Fig. 3, B) and more prominent in L5/6 (Fig. 3, C). By PM 9, the density of PV+ cell profiles had noticeably increased (Fig. 3, D, and E), and neuropil labeling was perpendicular to the surface.

**Figure 3:**
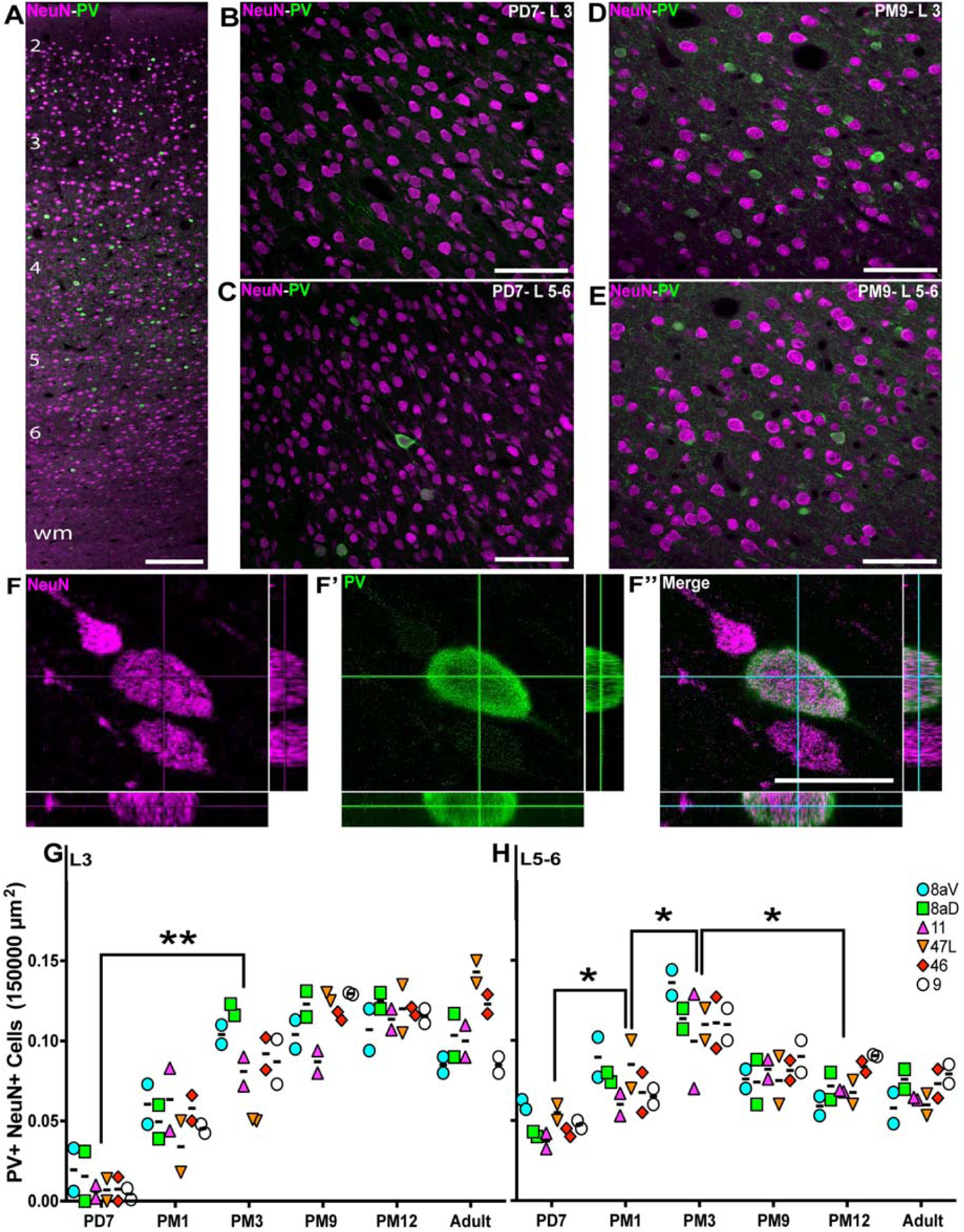
Progressive increase of the proportion of PV+ interneurons in the marmoset PFC from birth to adolescence. **A** PV+ interneurons (green) and NeuN+ neurons (magenta) are present across all cortical layers the marmoset DLPFC (area 46) at PD7. Distribution of PV+ NeuN+ interneurons in area 46 L3 and 5/6 of at PD7 (**B**, **C**) and PM9 (**D**, **E**). **F-F”** Cutlines through a 3D stack acquired with a laser scanning confocal microscope confirm that NeuN nuclear labeling and cytoplasmic PV labeling correspond to a unique cell in area 46 L3 at PD7. The ratio of PV+ NeuN+ interneurons over the total number of NeuN+ neurons per 150,000µm^2^ was calculated at each developmental stage in all 6 PFC areas of interest and plotted over time. The proportion of SST+ interneurons increases progressively and at a comparable rate in all areas in the supragranular L3 (**G**). In the L5/6 (**H**), the proportion of PV+ NeuN+ initially increased, like L3, followed by a reduction at the start of adolescence. Statistical analysis was performed using a nonparametric Kruskal–Wallis test followed by Dunn’s multiple comparisons. The data are presented as median ± interquartile ranges, * p<0.05: significant (n=2).

We confirmed that PV+ INs expressed the pan-neuronal transcription factor, NeuN (Fig. 3, F-F”), and proceeded to quantify the ratio of PV+ INs. The results in L3 confirmed our initial observations, revealing very few PV+ cells at PD7 across all PFC areas observed (0.003 ± 0.0007; Fig. 3, G). The proportion increases rapidly over the next three months (0.094 ± 0.06, p≤ 0.005, Kruskal-Wallis test), stabilizing to adult levels around 9 months old.

For L5/6, the average proportion of PV+ INs was greater at PD7 (Fig. 3, H; 0.045 ± 0.02). It increased over the first 3 months of life, peaking at PM3 (0.12 ± 0.03, p≤0.02, Kruskal– Wallis test), followed by a reduction over adolescence (0.07 ± 0.03, p≤0.02, Kruskal–Wallis test), then remaining constant into adulthood (0.07 ± 0.02). While deep cortical layers exhibited a distinct profile from superficial L3, all PFC areas revealed a similar profile in distribution, suggesting a common regulatory mechanism. These analyses also revealed that the fraction of PV+ INs is more significant than that of SST+ across all developmental stages observed.

Establishing a functioning network of PV+ INs is not limited to the number of INs, which does not tend to vary between healthy and diseased brains in the context of SCZ or ASD [46, 47], but also the level of PV expression, which can make these INs vulnerable to stressors. To assess expression levels, we calculated the intensity of the PV signal across all cortical layers. To account for background noise, values were normalized to the intensity of the signal in L1, devoid of PV expression. For example, the signal in area 46 was exclusively located in L5–6 at PD7 (Fig. 4, A), progressively expanding in L4, 3, and 2 over the first 3 months of life. The signal intensity, comprising cell bodies and neuropil immunostaining, was greatest during adolescence (PM9–12), remaining lighter in L2 than in all the other layers. In the adult, the intensity remained strong in L3 and 4 but dropped in L5–6, corresponding with the reduction of cell body count described above.

**Figure 4:**
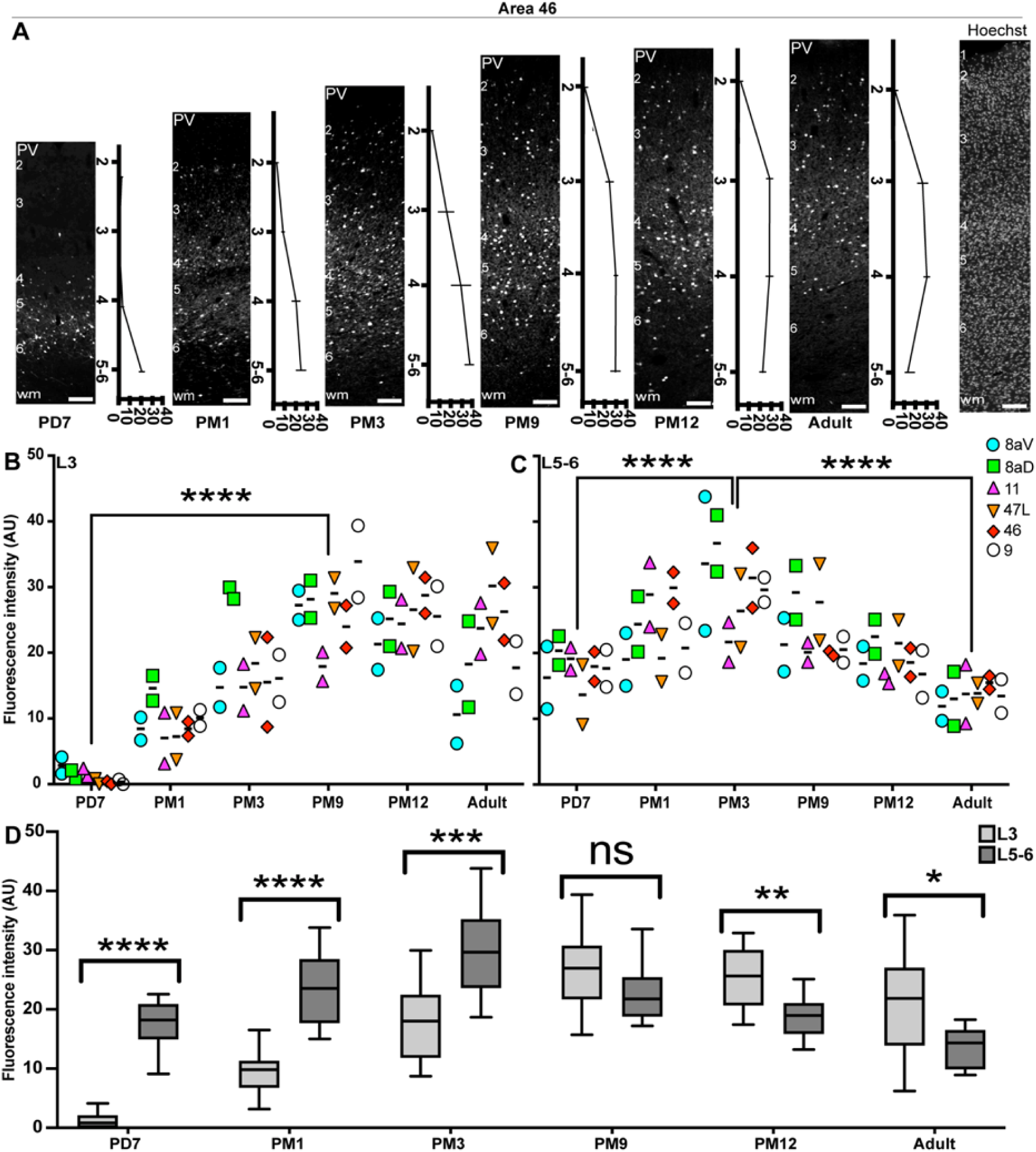
Cellular and neuropil expression of PV consistently increases from birth to adolescence across the marmoset PFC. **A** Laminar distribution of PV immunofluorescence (normalized to L1) across different ages in area 46. Hoechst labeling was used to demarcate the individual cortical layers. **B** and **C** Quantification of PV signal intensity revealed a consistent increase in PFC L3 and 5–6 from birth to adolescence. While levels remained stable in layers during adulthood, they decreased in L5/6 from PM9 to adult. However, PV FI significantly decreased from PM3 until adulthood. **D** Comparison of the PV immunofluorescence signal intensity in L3 and 5–6 in areas 8aD, 8aV, 47, 11, 9, and 46. Statistical analysis was performed using a nonparametric Mann–Whitney test. Data presented as median ± interquartile ranges (n=2), * p<0.05: significant, ns: non-significant.

We applied this method to all areas of interest in the PFC and plotted the average value for L3 (Fig. 4, B, S2) and 5–6 (Fig. 4, C, S2). The results follow the same profile as the cell counts, suggesting a correlation between cell counts and PV intensity across the developmental stages examined.

We then compared the intensity of the PV signal between L3 and L5–6 for each developmental stage (Fig. 4, D), confirming that the fluorescence in L3 was consistently lower until the beginning of adolescence (PM9), at which point an inflection occurred and L3 became predominant. This suggests that adolescence is a turning point in establishing adult PV function.

### Consolidation of PV+ interneuron connectivity in the PFC occurs during adolescence

The ultimate step of cortical maturation corresponds to the consolidation of the synaptic connections that persist beyond the phase of synaptic pruning [48]. Consolidation consists of two significant events: axonal myelination [49] and the deposition of chondroitin sulfate proteoglycans (CSPG), one of the main components of perineuronal nets (PNN), which form the extracellular matrix (ECM) [50]. The CSPG scaffold encapsulates the pre-and postsynaptic elements, creating a mesh-like structure stabilizing the synapse [51] aggregation of CSPG, which occurs late in development. It correlates with establishing mature synapses following a ‘critical period.’ PNNs are heterogeneously enriched across the neocortex, their distribution profile enabling the parcellation of discrete cortical areas [53]. To assess the level of CSPG scaffold, PV+ synapses in the marmoset PFC, we used *Wisteria floribunda Agglutinin* (WFA) labeling to reveal the accumulation of CSPG in PNN throughout postnatal development and into adulthood. For area 46, WFA cell-profile labeling was observed predominantly in L3–5 and the white matter in adulthood (Fig. 5, A). At PM1, faint WFA labeling could be observed in L3 but did not correlate with PV+ INs (Fig. 5, B). However, in L5–6, WFA colocalized with PV labeling (Fig. 5, C). At 12 months old, most PV+ INs were outlined by WFA labeling, including their proximal neurites in L3 (Fig. 5, D) and L5–6 (Fig. 5, E). High-magnification confocal images of PV-WFA double-positive INs illustrate how PNN thickly envelops the cell body and the extensive proximal neurites (Fig 5, F-F”).

**Figure 5:**
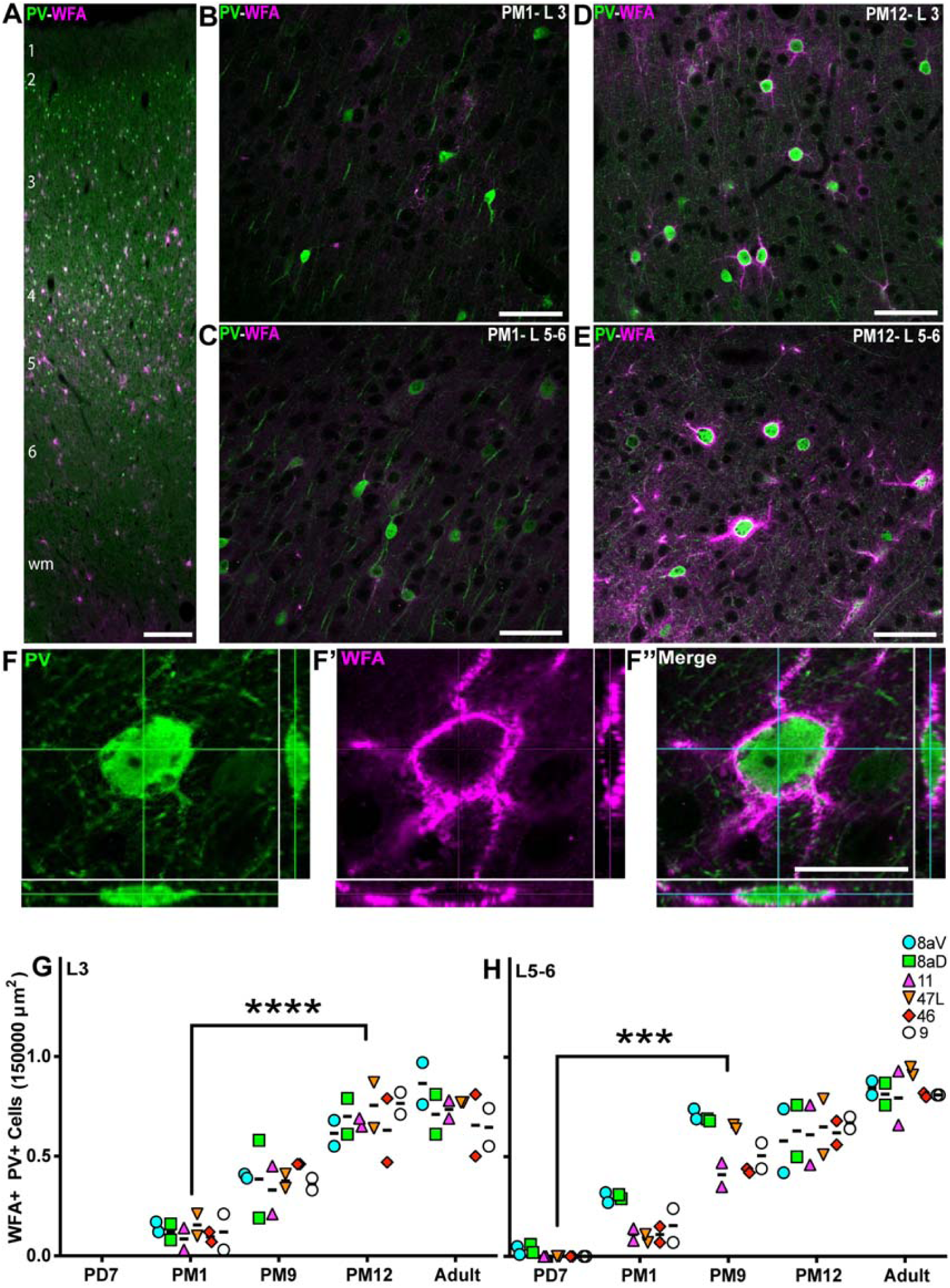
Progressive accumulation of WFA+ perineuronal nets on PV+ dendritic tree from birth to adult across the marmoset PFC. **A** PV+ interneurons (green) and WFA+ membrane (magenta) are present across all cortical layers the marmoset DLPFC (area 46) at PD7. Distribution of WFA+ PV+ interneurons in area 46 L3 and 5/6 of at PM1 (**B**, **C**) and PM12 (**D**, **E**). **F-F”** Cutlines through a 3D stack acquired with a laser scanning confocal microscope confirm that PV cytoplasmic labeling and membrane-bound WFA labeling correspond to a unique cell in area 46 L3 at PM12. The ratio of WFA+ PV+ interneurons over the total number of PV+ neurons per 150,000µm^2^ was calculated at each developmental stage in all 6 PFC areas of interest and plotted over time. The proportion of WFA+ interneurons increases progressively and at a comparable rate in all areas in the supragranular L3 (**G**) and infragranular L5/6 (**H**). Statistical analysis was performed using a nonparametric Kruskal–Wallis test followed by Dunn’s multiple comparisons. The data are presented as median ± interquartile ranges, * p<0.05: significant (n=2).

To support these observations, we further quantified the fraction of PV+ INs cloaked with WFA. In L3, the PV+ and WFA+ cells were observed at PM1. The proportion of double-positive PV+ INs steadily increased over development, reaching a plateau by PM12, when over 80% of PV+ INs exhibited PNN, consistent across all PFC areas of interest (Fig. 5, G). The time course of PNN accumulation on PV+ INs in L5/6 preceded that of L3 (Fig. 5, H) as the first WFA+ PV+ double-positive INs were observed as early as PD7 but otherwise followed a comparable time course, peaking between PM12 and adulthood. Interestingly, areas 8aV and 8aD revealed an earlier peak of WFA+ PV+ INs in L5/6 at PM9 compared to other PFC areas.

Additionally, the precise changes in fluorescence intensity of WFA around the cell membrane of PV+ cells were investigated in three prefrontal areas of 11, 47L, and 46 (S3, A, and B). The data analysis revealed a progressive increase in WFA fluorescence intensity from PM1, reaching an adult-like level in PM12 (21.6 ± 16.6 vs. 380.8 ± 40, p=0.01) (S3, A). Likewise, in layers 5-6, the fluorescence intensity of WFA significantly increased from PD7 to PM12 (2.6 ± 4.3 vs. 524 ± 155, p=0.01). A further 64% increase was detected from PM12 until adulthood. However, it was not identified to be statistically significant (524 ± 155 vs. 861.5 ± 222, p>0.9) (S 3, B). These findings are consistent with cell counting data indicating that the accumulation of PNNs around PV+ cells was a protracted process until mid-adolescence.

### Upregulation of KCC2 in the cell membrane of PV+ interneurons in infancy

The upregulation of PV in young INs is believed to occur through activity-dependent pathways [54] to increase Ca^+2^ buffering capability, leading to burst firing [55]. For this reason, PV is a reliable indicator of IN maturation. However, PV is not unique, and other cellular aspects can predict functional maturation. In particular, the potassium chloride co-transporter, KCC2. For example, as PV interneurons mature, there is a significant upregulation of KCC2. This increase in KCC2 expression lowers the intracellular chloride concentration, allowing GABAergic inputs to become hyperpolarizing and thus inhibitory. [26, 56].

To determine the extent to which KCC2 informs the maturation of INs during the postnatal development of the marmoset PFC, we analyzed the expression of KCC2 on PV+ INs. At PD7, we observed the presence of KCC2 puncta on the putative cell membrane of PV+ cell profiles in L5/6 (Fig. 6, A), which was more uniform and ubiquitous than PM1 (Fig. 6, B). To confirm this observation, we quantified the intensity of the KCC2 signal along the outline of PV+ cell bodies (Fig. 6, C-C”), as previously described [57].

**Figure 6:**
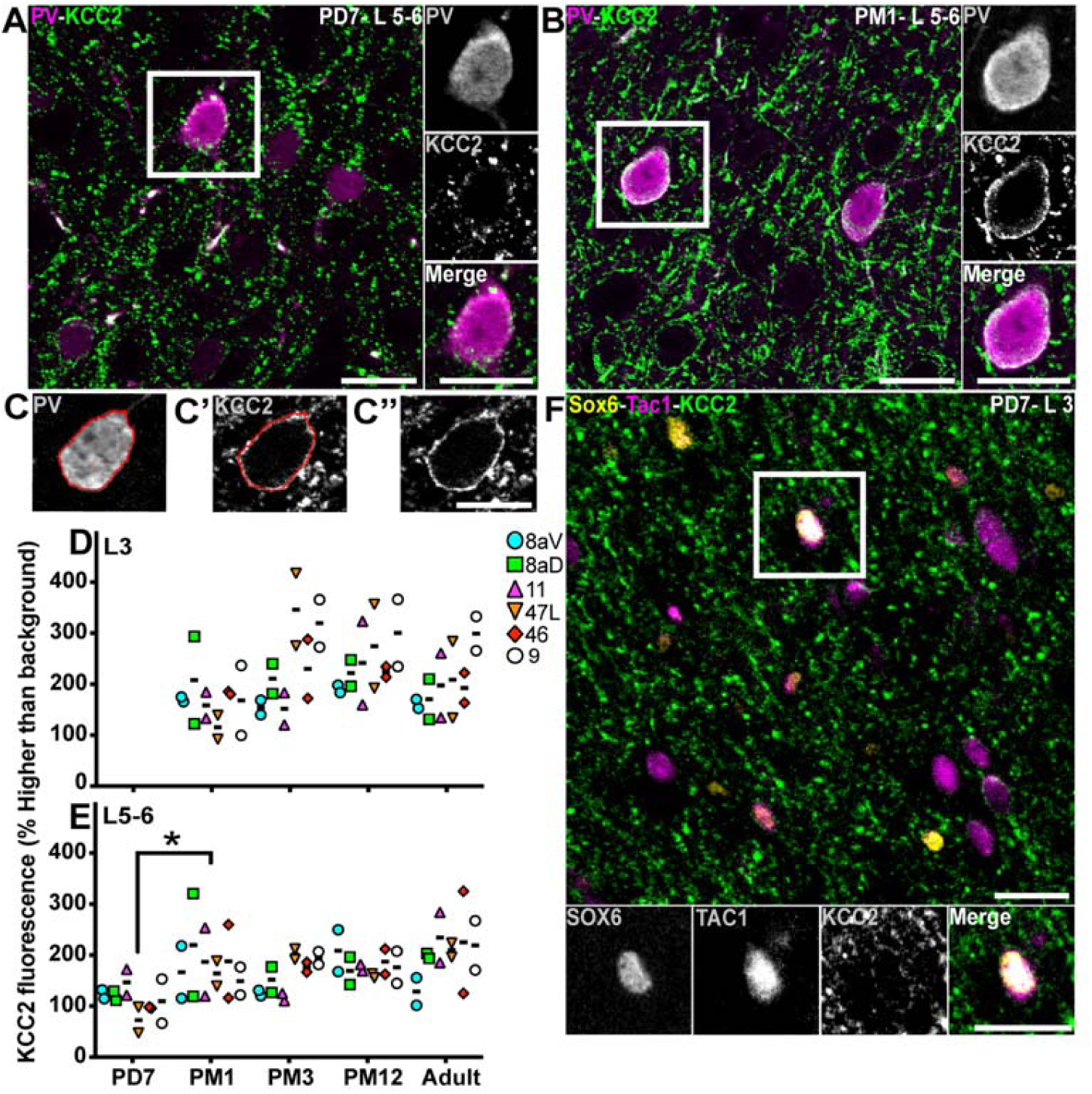
Expression of KCC2 in PV+ cells and prospective PV+ interneurons from birth to adulthood in the marmoset PFC. PV (magenta) and KCC2 (green) labeling in L5–6 of area 46 at PD7 (**A**) and PM1 (**B**) Representative example of the quantification of KCC2 fluorescence intensity on the membrane of PV+ interneuron with first, tracing the outline of the PV+ cytoplasm (**C**), pasting this outline on the channel corresponding to KCC2 labeling (**C’**) to measure the fluorescent signal intensity and the same image without the outline (**C”**) The intensity of the KCC2 fluorescent signal intensity on PV+ interneurons of the marmoset PFC L3 (**D**) and 5–6 (**E**) was plotted across development. **F** Presumptive PV+ interneurons identified by Sox6 (yellow) and Tac1 (magenta) express KCC2 (green) in L3 of area 8a in neonates (PD7), suggesting electrical activity of precursors of PV+ cells before PV upregulation. Statistical analysis was performed using the nonparametric Kruskal–Wallis test, followed by Dunn’s multiple comparisons. Data presented as median ± interquartile ranges, * p<0.05: significant (n=2).

In L3, the amount of KCC2 signal around PV+ INs did not vary across development or areas (Fig. 6, D). In L5–6, the amount of KCC2 signal on PV+ INs increased twofold between PD7 and PM1 (Fig. 6, E), where it remained constant into adulthood.

The steady KCC2 signal around PV+ INs over development in L3 suggests that KCC2 upregulation might precede that of PV. To test this hypothesis, we identified a combination of proteins to label a presumptive subgroup of PV+ INs, namely Sox6 and TAC1. Transcription factor Sox6 is expressed in all INs emerging from the MGE [58]. At the same time, TAC1 is a neuropeptide notably involved in increasing cytosolic Ca^+2^ ions [59] and is expressed in various neuronal fractions, including PV+ INs [60]. We randomly selected area 8aD, tested the hypothesis in L3 at PD7 (when there are few PV+ cells in L3; see Fig 3), and identified distinctive KCC2+ puncta outlining the surface of putative PV+ INs (Fig. 6, F). This suggests that KCC2 expression precedes that of PV.

## Peak expression of fast-spiking phenotype in PV+ Interneurons during postnatal development

Mature cortical PV+ INs generate fast-spiking action potentials. This characteristic is critical for the normal function of the adult neocortex. The potassium channel, Kv3.1b, and sodium channel, Nav1.1, are essential for generating fast-spiking activity in PV+ INs [25]. To estimate the postnatal stage at which PV+ INs have the capacity for fast-spiking action potentials, we used the same analytical approach as for KCC2 to analyze the distribution of Kv3.1b and Nav1.1 on PV+ INs in the maturing PFC. Kv3.1b was expressed at the surface of PV+ INs in L3 as early as one month old (Fig. 7, A) and persisted at 3 months (Fig. 7, B). We also observed Nav1.1 expression around PV+ INs at PM1 (Fig. 7, C) and PM3 (Fig. 7, D).

**Figure 7:**
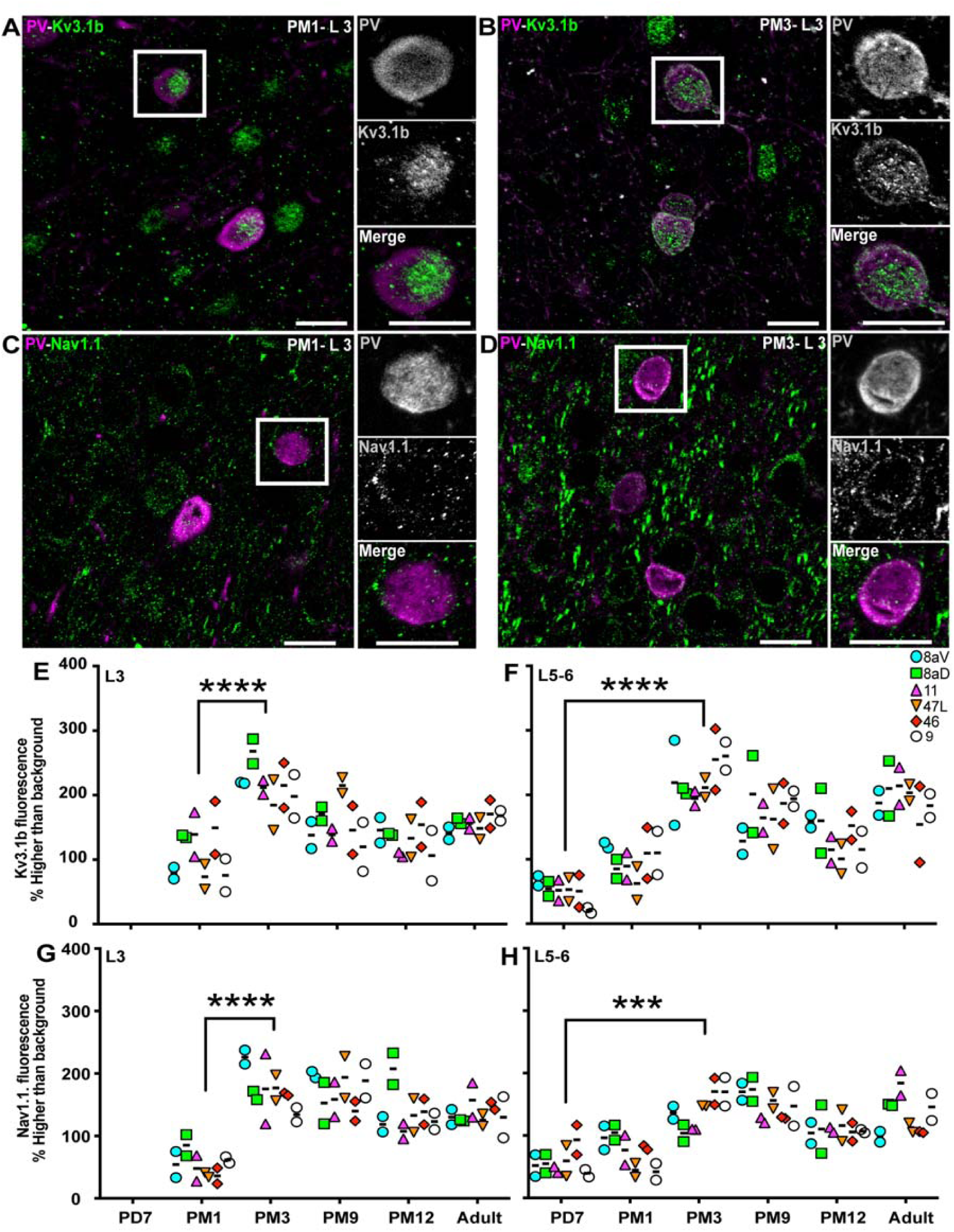
Expression Kv31.b and Nav1.1 ion channels progressively increase on the membrane of PV+ neurons across the marmoset developing PFC. Correlation of Kv3.1b (green) expression and PV (magenta) in interneurons in L3 of area 46 at PM1 (**A**) and PM3 (**B**) Correlation of Nav1.1 (magenta) expression and PV (green) in interneurons in area 46 L3 at PM1 (**C**) and PM3 (**D**) Intensity of Kv3.1b fluorescent signal in PV+ interneurons in L3 (**E**) and 5–6 (**F**) of the marmoset PFC plotted over time reveals an increase during the first 3 months postnatal Intensity of Nav1.1 fluorescent signal in PV+ interneurons in L3 (**G**) and 5–6 (**H**) of the marmoset PFC plotted over time reveals an increase during the first 3 months postnatal. Statistical analysis was performed using the nonparametric Kruskal–Wallis test, followed by Dunn’s multiple comparisons. Data presented as median ± interquartile ranges, * p<0.05: significant (n=2).

The quantitative analysis across all areas of interest revealed that although Kv3.1b was not detected on PV+ INs at PD7 in L3, the amount of Kv3.1b on PV+ interneurons doubled between PM1 and PM3 (Fig. 7, E). Kv3.1b signal appeared to decrease during adolescence. In L5/6, Kv3.1b signal was present around PV+ INs from PD7 onwards, increasing steadily over the first 3 postnatal months and remaining constant throughout adolescence and adulthood (Fig. 7, F).

The profile of Nav1.1 expression was comparable to that of Kv3.1b, with no signal observed at PD7 in L3 and a doubling of the intensity of the signal between PM1 and PM3 (Fig. 7, G). The expression remained constant from PM3 to adulthood. In L5–6, Nav1.1 was detected around PV+ INs from PD7, increasing over the first 3 postnatal months (Fig. 7, H) to a lesser extent than Kv3.1b.

The analysis of transcriptomics and electrophysiological data has previously demonstrated a strong positive correlation between 1) high expression of Kv3.1b and 2) a balanced index calculated by Kv3.1b/Nav1.1 ratio, which determines the fast-spiking phenotype of PV+ INs [25]. To predict at what age during postnatal development the fast-spiking phenotype of PV+ cells may be the highest, we applied the Kendall correlation test between the Kv3.1b expression level and the Kv3.1/Nav1.1 ratio in L3 and 5–6 between PM3 and adulthood (Fig. 8, A-D). In L3, the Kendall test revealed the highest correlation coefficient at PM9 (coefficient value=0.63) (Fig. 8, B and E). The correlation coefficient decreased 0.43-fold lower in adulthood than PM9 (Fig. 8, B, D, and E). In L5–6, significant correlation coefficients were also detected during adolescence, PM9 and PM12 (coefficient values of 0.77 and 0.89, respectively) (Fig. 8, B, C, and F), and a 0.66-fold decrease was evident in adulthood compared with PM12 (Fig. 8, C, D and F).

**Figure 8:**
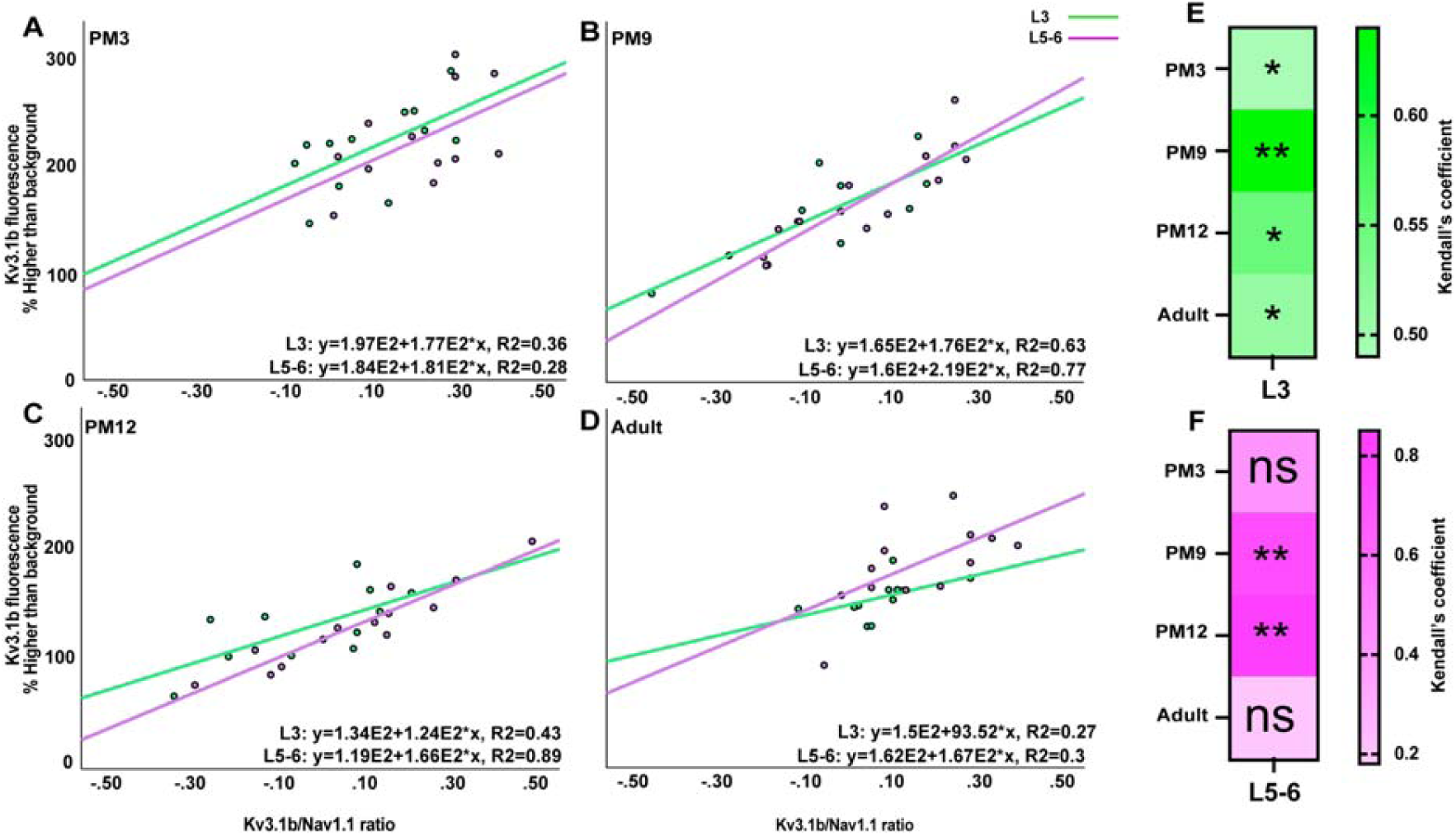
Highest correlations between factors essential for fast-spiking properties of PV+ interneurons during adolescence. Kendall’s correlation between the fluorescence intensity of Kv3.1b and ratio of Kv3.1b to Nav1.1 in L3 (green) and L5–6 (magenta) of PFC at PM3 (**A**), PM9 (**B**), PM12 (**C**) and adulthood (**D**), at which the highest correlation was observed (see Fig. 7) The heat map highlights PM9 as the age during which Kendall’s correlation coefficient was the highest consistently in L3 (**E**) and 5–6 (**F**), suggesting the maturation of the fast-spiking phenotype of PV+ interneurons occurs during adolescence. * p<0.05: significant (n=2).

## Discussion

This study scrutinized the postnatal maturation trajectory of the MGE-derived IN network, encompassing PV and SST subtypes, within the PFC of the marmoset monkey. The PFC’s intricate developmental and maturation processes entail a multifaceted interplay of diverse cellular and molecular mechanisms, hypothesized to undergo a temporal delay relative to maturation processes in the sensorimotor cortex. The failure of several molecular and cellular processes in PFC has been associated with several neurodevelopmental disorders [61–67]. INs are crucial in maintaining the brain’s neural activity balance, regulating the timing and synchronization of neural signals, and ensuring fine-tuning of the overall functioning of the neural circuitry [68]. Interestingly, we observed a similar maturation profile across the distinct PFC areas (8aD,8aV, 9, 11,46, and 47) investigated. However, the maturation profile for SST and PV revealed a converse relationship from infancy into adulthood, whereby SST neurons peaked by PD7, yet PV neurons only peaked by adolescence (PM9). Considering the protracted developmental profile of PV neurons in the PFC and their contribution to higher cognitive functions [46, 69–78], we elucidated the spatiotemporal upregulation of three ion channels, namely KCC2, Kv3.1b, and Nav1.1, which are intricately associated with distinct functional aspects of PV+ INs.

### Unraveling the discrepancy in maturation of prefrontal cortex SST interneurons: primate vs. rodent

The developmental profile of SST INs in PFC is still debated and may be species-specific. We revealed in the marmoset PFC that the fraction of SST INs is highest in the first postnatal week and steadily decreases until adolescence, after which it stabilizes. This is consistent with human microarray data, which revealed a gradual postnatal decrease in SST mRNA in the human PFC [69], whereas the number remains stable in the mouse medial PFC over the first 40 days of life [79]. Our results are consistent with the human data, suggesting a divergence between primate and rodent SST INs in the PFC. A recent comparative analysis of the transcript signature of the rodent, non-human primate, and human PFC confirms this conclusion, demonstrating Order-specific modification of interneuronal composition and a more significant proportion of cortical INs in primates compared to rodents (25–34% vs. 15– 20%) [6].

The alterations observed, coupled with the evolutionary enlargement of the PFC, incorporating newly identified regions, many featuring a granule cell layer, provide substantive evidence endorsing our hypothesis that the developmental progression of inhibitory neurons (INs) within the primate PFC postnatally follows distinctive mechanisms compared to rodents. These findings bear significant implications for selecting appropriate animal models in the exploration of neurodevelopmental disorders, such as schizophrenia, autism spectrum disorders, and ADHD.

### Accelerated maturation of SST interneurons compared to PV interneurons in the monkey PFC

The rapid decline in SST INs could imply SST functionally influences early neuronal circuit processes. SST is a multifunctional hormone previously linked to migration [80], regulation of proliferation [81], apoptosis [82], differentiation [81], synaptogenesis, and axon pathfinding [83] in maturing neuronal circuits. These functions are critical, particularly during early postnatal events of development. While it is not studied in primates, in the mouse somatosensory cortex, it was reported that SST INs strongly innervate PV+ INs at P6 in L5/6 and act as placeholders for establishing thalamocortical connectivity in the PV+ INs [45, 84]. This finding supports the earlier upregulation and maturation of SST+ INs compared with PV+ INs by mediating the establishment of thalamocortical inputs onto PV cells [85]. Consistently, evidence from *in situ* hybridization experimentations in the marmoset also indicates widespread SST mRNA present across the laminae of the PFC at birth (PD0); however, no PV mRNA was detected at the same prefrontal area at PD0 [86, 87]. A study in human DLPFC demonstrated a significant upregulation of PV mRNA between birth and juvenility, compared to a decreasing trend of SST mRNA [69]. This suggests that SST INs appear before PV+ INs in primates, including humans.

Interestingly, while there are apparent differences between primates and rodents, the upregulation of PV around adolescence has also been observed in the medial PFC of rats [14]. This corroborates the suggestion that PV is essential for the refinement of prefrontal GABAergic function and connectivity, during which PV upregulation supports the acquisition of the mature GABAergic phenotype necessary to sustain adult PFC functions.

### The maturation of the PV IN network is synchronized across the cortices of the PFC

Our analysis of the PV IN subgroup, the largest IN fraction in the primate neocortex, revealed the maturation of these cells is also synchronized across the PFC areas. These findings agree with a similar analysis of the maturation of NNF+ pyramidal neurons [30], with which the PV neurons extensively synapse, and again highlight the role of the PV IN network in maturation. Of note, previous studies in the primary visual cortex (V1) of marmoset and macaque monkeys reveal an adult-like level of PV+ INs was observed as early as the neonatal stage (PD14 and PD22, respectively) [29, 88]. This contrasts with the findings in this study, indicating a protracted maturation of PV+ INs in the PFC during early adolescence, suggesting spatiotemporal differences in PV expression profile among non-human primates’ cortex.

The main factor determining the timing of maturation appears to be the IN laminar identity, as we consistently observed that the infragranular layer exhibited evidence of maturation before their supragranular counterparts. For example, PV+ INs are present in L5/6 by the first week postnatal, but the presence in L3 is delayed. Similarly, the deposition of PNN on PV+ INs peaked by PM9 in L5/6 but only in PM12 in L3. This temporal profile matches NNF+ pyramidal neurons across the marmoset PFC and supragranular vs infragranular dichotomy [30].

Interestingly, not all PV+ INs exhibited PNN labeling in adulthood, suggesting that a proportion may remain amenable to synaptic remodeling into adulthood. Previous data indicate the neurons in adult PFC can modify their connectivity in response to changes in environmental factors such as stressors [89–91]. In the mouse visual cortex, it has been shown that PV+ cells continue to regulate plasticity in adulthood [92] , and manipulations of PNNs in adult PV+ cells have been identified to reinstate plasticity in the mouse visual cortex [93]. These findings support that the modifications of PNN deposition can be a mechanism by which PV cells+ regulate plasticity across postnatal development and adulthood. Another possibility is that not all PV+ subtypes express the components of PNNs. Moreover, as described by Ariza et al., among PV-expressing cell types, only basket cells are accumulated by PNN in the human PFC [78]. They reported that PNN labeling in PV+ cells provide tools for segregating the basket cells from other cell types expressing PV, such as chandelier cells [78].

### Ion channel expression suggests that the fast-spiking phenotype of PV interneurons may be the highest in adolescent PFC

We observed that KCC2 expression preceded that of PV and was associated with putative PV+ INs in the first week of postnatal development, followed by PV+ INs into adulthood. This early expression of the KCC2 protein suggests that the IN subtype already has the capacity for hyperpolarizing GABA_A_ receptor currents, and an increase in PV protein expression has been associated with an increase in KCC2 function during development. Importantly, KCC2 dysfunction and dysregulation of Cl^−^-homeostasis occur in neurodevelopmental disorders, including Down syndrome [94], fragile X syndrome [95], Rett syndrome [96, 97], and schizophrenia [98].

The main hallmark of PV+ INs lies in their ability to generate fast-spiking action potentials, a property regulated by high Kv31b ion channel expression and a balance between Kv3.1b and Nav1.1 ion channels [25]. Among INs, the potassium channel transporter, Kv3.1b, is only expressed in PV+ INs across rodents and primates [99]. Mounting evidence indicates that the Kv3.1b ion channel plays a critical role in the rapid repolarisation of the PV IN membrane following a sodium channel-mediated depolarization [56, 100, 101]. Therefore, this ion channel is a central regulator of neuronal activity in PV+ INs. In addition to Kv3.1b, a sodium channel, Nav1.1 ion channel, is also demonstrated to be highly expressed on PV+ INs in the cell membrane and axon initial segment [25]. It contributes to the rapid depolarisation of the membrane and action potential propagation. As such, the expression of the Nav1.1 ion channel is representative of the neuronal activity within PV cells. Our findings demonstrate an augmentation in the fast-spiking potential of PV+ INs across the primate PFC’s supra- and infragranular layers during early adolescence. Significant remodeling and refinements of primate PFC networks have been postulated to occur at this stage, followed by consolidation of pathways terminating their development in adolescence [102–104]. Enhancement of the fast-spiking phenotype of PV+ INs during adolescence suggests their involvement in these developmental processes occurring in PFC at this age. To support this hypothesis, it has been previously demonstrated that inhibiting PV cell activity during adolescence has led to abnormal development of the frontal neocortex in rodents and subsequent cognitive impairments in adult animals [105]. The fast-spiking functions of PV cells are likely associated with inhibiting weak connections, thereby facilitating immuno-system-mediated synaptic pruning during adolescence. Also, at this age, the PFC receives extensive excitatory thalamic inputs related to its development, including the medial pulvinar and medial dorsal (MD) nucleus [4]. Further, the MD thalamocortical input to mPFC in mice was demonstrated to be crucial for the normal development of mPFC during adolescence, in which its disruption had a profound negative functional effect [106].

Consistent with these results, it has been shown that in rodents, the fast-spiking firing of PV+ INs experience a significant enhancement during adolescence (PD 25) in correlation with the rise in the gamma oscillations [107–109]. Robust gamma activity, mediated by the fast-spiking phenotype, has also been observed during primate adolescence in the PFC [110]. Given the contribution of gamma oscillations in mediating cognitive functions such as working memory in this region [111, 112], it could be presumed that adolescence is involved in the critical period of PFC maturation [104, 113]. Consistently, the external and internal factors that are associated with the onset and closure of critical period (e.g., brain-derived neurotrophic factor (BDNF), orthodenticlehomeobox2 (Otx2), and PNN) are known to contribute to PV+ cell maturation [114, 115]. Thus, the enhancement of PV+, the fast-spiking phenotype of PV+ cells during adolescence, as indicated in this research, reinforces the notion that these cells play a role in mediating the critical period of PFC development during this stage [116, 117].

### Statistical power vs temporal resolution

Our findings reinforce the discrepancy between rodents and primates regarding the brain’s functional and anatomical organization and cellular composition. This is particularly the case when investigating the PFC, for which most areas and functions have no equivalent in rodents. While research in the developing nonhuman primate offers a translatability that rodents cannot afford, it comes with essential drawbacks, principally the cost and availability of individuals. For this study, we privileged the temporal resolution, using 6 developmental stages over the statistical power (n=2) for 12 animals. Fortunately, the lack of inter-areal variation allowed us to cluster the different areas together and identify trends throughout development. Like previous research [30], the small sample size limits our ability to speculate. Still, the consistency across our observations and others consolidates our conclusion that PFC areas mature uniformly and not sequentially. We also provide novel insights on various aspects of PV IN physiology, including ion channel expression and PNN deposition.

## Materials and methods

### Animals

Twelve marmoset monkeys (*Callithrix jacchus*) aged postnatal day (PD) 7 (n=2; 0 male: 2 female); postnatal month (PM) 1 (n=2; 1:1); PM3 (n=2; 1:1); PM9 (n=2; 0:2); PM12 (n=2; 1:1); and adult (>2.5years; n=2; 1:1) were selected for this study. Gender was not a criterion in the selection of animals. Animals were procured from the National Non-human Primate Breeding and Research Facility (Australia) and housed in a vivarium (12:12 hr light/dark cycle, temperature 31°C, humidity 65%). All experiments were conducted in accordance with the Australian Code of Practice for the Care and Use of Animals for Scientific Purposes and were approved by the Monash University Animal Ethics Committee, which also monitored the welfare of the animals.

### Tissue processing

Animals were administered an overdose of pentobarbitone sodium (100 mg/kg). Following apnoea, neonates and juveniles were transcardially flushed with warm (∼30C) heparinized phosphate buffer 0.1M (PB; pH 7.2) containing 0.1% sodium nitrite and adults with room temperature heparinized saline (0.9%). All animals were subsequently perfused with 4% paraformaldehyde in PB 0.1M and postfixed overnight in 4% paraformaldehyde at 4°C, dehydrated in increasing concentrations of sucrose (10, 20, and 30%) in PB 0.1M, frozen over a bath of liquid nitrogen, and stored at −80°C until cryosectioning.

### Histology and immunolabelling

Each brain hemisphere was cut in the coronal plane on a cryostat (CM3050S; Leica, Wetzlar, Germany) at a thickness of 50µm, divided into four series, and stored free-floating in a cryoprotective solution (50% phosphate buffer saline 0.1M PB, 30% ethylene glycol, 20% glycerol) at −20°C. Sections were rinsed in PBS and blocked in a solution of PBS, 0.3% Triton-X, and 5% normal donkey serum for 1 hour at room temperature. Sections were incubated with the primary antibodies (combined in the case of double-labeling) and biotin-conjugated WFA (1:400, L1516, Sigma) for PNN labeling, diluted in the blocking solution (detailed in Table 1) for 16–18 hours at 4°C. For SST/NeuN double-labeling, Triton-X was omitted from the blocking solution, and sections were incubated for 48–72 hours. Following incubation, sections were rinsed three times in PBS for 10 min each, incubated with the appropriate secondary antibodies in donkey anti-rabbit Alexa Fluor 488 (Thermo, USA, A10039; 1:800), donkey anti-guinea pig Alexa Fluor 488 secondary antibody (Abcam, USA, ab150185), donkey anti-mouse Alexa Fluor 594 (Thermo, A21203; 1:800), donkey anti-mouse Alexa Fluor 488 secondary antibody (Thermo, A21202; 1:800), donkey anti-rabbit Alexa Fluor 594 secondary antibody (Thermo, A21207; 1:800), and goat anti-rat Alexa Fluor 488 secondary (Thermo, A11006; 1:800) antibodies as well as streptavidin Alexa Fluor 594 (Thermo, A511227; 1:800) (Molecular Probes, Thermo, La Jolla, CA) in the blocking solution for 1 hour, rinsed three times in PBS for 10 min, incubated with Hoechst (Pentahydrate bis-Benzimide, Dako, cat# H1398) to visualize cell nuclei. The sections were then rinsed in PBS, mounted on Superfrost Plus glass slides (Thermo) with Fluoromount G mounting medium (Thermo), and coverslipped. The labeling was carried out only with a secondary antibody for negative control.

**Table 1:** list of primary antibodies employed in the study.

| Antigen | Abbreviation | Host Species | Dilution | Source | Catalog/clone |
| --- | --- | --- | --- | --- | --- |
| Neuronal nuclear antigen | NeuN | Rabbit | 1:1000 | Merck | MAB377 |
| Parvalbumin | PV | Mouse and Rabbit | 1:1000 | Swant | PV28, PV235 |
| Somatostatin | SST | Rat | 1:100 | Merck | MAB2358 |
| Active Caspase 3 | aCasp3 | Rabbit | 1:200 | Thermo Fisher | MA5-32015 |
| Potassium/Chloride cotransporter 2 | KCC2 | Rabbit | 1:150 | Cell Signalling Technology | 94725 |
| Substance p | Tac1 | Guinea pig | 1:500 | Boster Bio | A06666 |
| Transcription factor | Sox6 | Mouse | 1:200 | Thermo Fisher | MA5-31426 |
| Potassium channel | Kv3.1b | Rabbit | 1:500 | Merck | AB5188 |
| Sodium channel | Nav1.1 | Rabbit | 1:200 | Alomone Labs | ASC-001 |

### Microscopy

Sections were imaged with an Axio.Imager Z1 epifluorescent microscope (Zeiss, Germany) equipped with an Apotome to acquire near-confocal quality Z-stacks. Three images of each section were obtained with a Zeiss Axiocam HRm digital camera using the Axiovision software (v 4.8.1.0) at a resolution of 1024L×L1024 pixels, saved as Zeiss Vision Image (ZVI) and exported to tagged image file format (TIFF) format. The objectives used were Zeiss EC-Plan-Neofluar 5L×L0.16, #420330-9901, EC-Plan-Neofluar 10L×L0.3, #420340- 9901, and Plan Apochromat 20L×L0.8, #420650-9901. Filter sets used for visualizing fluorescently labeled cells were Zeiss 49 4’,6-diamidino-2-phenylindole (DAPI) #488049- 9901-000, Zeiss HE enhanced Green Fluorescent Protein (eGFP) #489038-9901-000, and Zeiss HQ Texas Red #000000-1114-462.

Low-magnification photomicrographs (1300 × 1030 dpi) were acquired with a Zeiss Discovery V20 stereomicroscope and an Axiocam HRc camera connected to Axiovision (version 4.7.1) (Zeiss Microscopy, LLC, NY, USA).

To confirm the colocalization of SST/NeuN, PV/NeuN, and WFA/NeuN double immunostainings, a Z-stack was captured on a C1 Invert Confocal Microscope (Nikon, Tokyo, Japan). The microscope was equipped with excitation laser lines at 405, 488, 561, and 638 nm, and the images were obtained at a scan size of 1024 × 1024 bpi, with frame averaging set to 2. Two objectives, X40 and X60, were used. The acquired images were processed using NIS software (Nikon, Tokyo, Japan) and analyzed in Fiji (NIH).

### Digital image processing and quantification

Image stitching, contrast, and brightness adjustments were performed using Adobe Photoshop CC 2019. Figures, including contours, labels, and annotations, were composed using Adobe Illustrator CC 2019.

For PV, SST, and PNN labeling quantification, only cortical L3 and L5/6 were considered. These layers were selected because they act as input and output in the PFC. Hence, their maturation reflects neuronal network connectivity maturation in the PFC. For each animal, three images of each cortical area were randomly captured, and cell quantification was performed using the Fiji *Cell Counter* plugin [34]. The counting frame was estimated to be 440 × 340 µ^2^. The results were shown as a ratio of SST+ NeuN+ cells, PV+ NeuN+ cells, and WFA+ PV+ cells. For each age, the fluorescence intensity of L3 and 5–6 was measured using Fiji image software [118], and the result was normalized to the fluorescence intensity of L1, which did not exhibit a signal and only emitted autofluorescence, often a result of aldehyde fixation. The ion channel fluorescence intensity was quantified in an average number of 20 cells by Fiji according to the previously described methods [119]. Briefly, the single plane images (with the objective of X40) including the largest surface area of PV+ cells were considered for ion channel analysis. The PV+ cell membrane (mode: 8-connected; tolerance: 1000) was outlined automatically using wand tracing tool in Fiji. The trace was converted to a line with 0.5 μm width, and the average intensity of the ion channel signal convergent with the line was calculated (grey level, 12 bits). The values obtained were normalized to the autofluorescence signal in the background, obtained from a 10µm^2^ square area in the same optical plane adjacent to the PV+ cells and lacking labeled neuropil. The results were presented as a percentage over the background using the following equation:

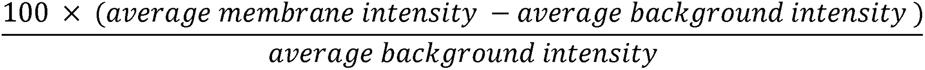

To predict the developmental stage at which PV+ INs exhibit fast-spiking action potential, the association between the fluorescence intensity of ion channels Kv3.1b and Nav1.1 was profiled across postnatal development using the ratio:

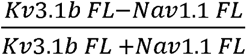

### Statistical analysis

Statistical analyses were performed using GraphPad Prism software version 9.5.1. All data are presented as median ± Interquartile range. A *p-*value of ≤ 0.05 was considered statistically significant. A nonparametric Kruskal–Wallis test followed by Dunn’s posthoc test was applied to the cell count ratio and ion channel analysis data. The data were presented as median ± interquartile ranges.

A Mann-Whitney test compared the fluorescence intensity levels between L3 and 5–6 for all stages. The data are presented as median ± interquartile ranges. A *p*-value <0.05 was considered significant. A nonparametric Kendall correlation test was performed to detect an association between Nav1.1 and Kv3.1b at the different developmental stages.

## Acknowledgments

The authors thank Monash MicroImaging core facilities for their contribution to this work and acknowledge the assistance of Monash Statistical Consulting Service for data analysis. This research was supported by an Australian Research Council grant (DP190101948) and the Intramural Research Program of the NIMH (ZIA MH002984) to J.A.B. The Australian Regenerative Medicine Institute is supported by grants from the State Government of Victoria and the Australian Government.

Data is available upon request.

