## Supplementary material for "Developmental dynamics of the prefrontal cortical SST and PV interneuron networks: Insights from the monkey highlight human-specific features": Figs S1-S3 (from another version of this preprint: version 1, 2024-07-10)

**Supplementary (S) data**

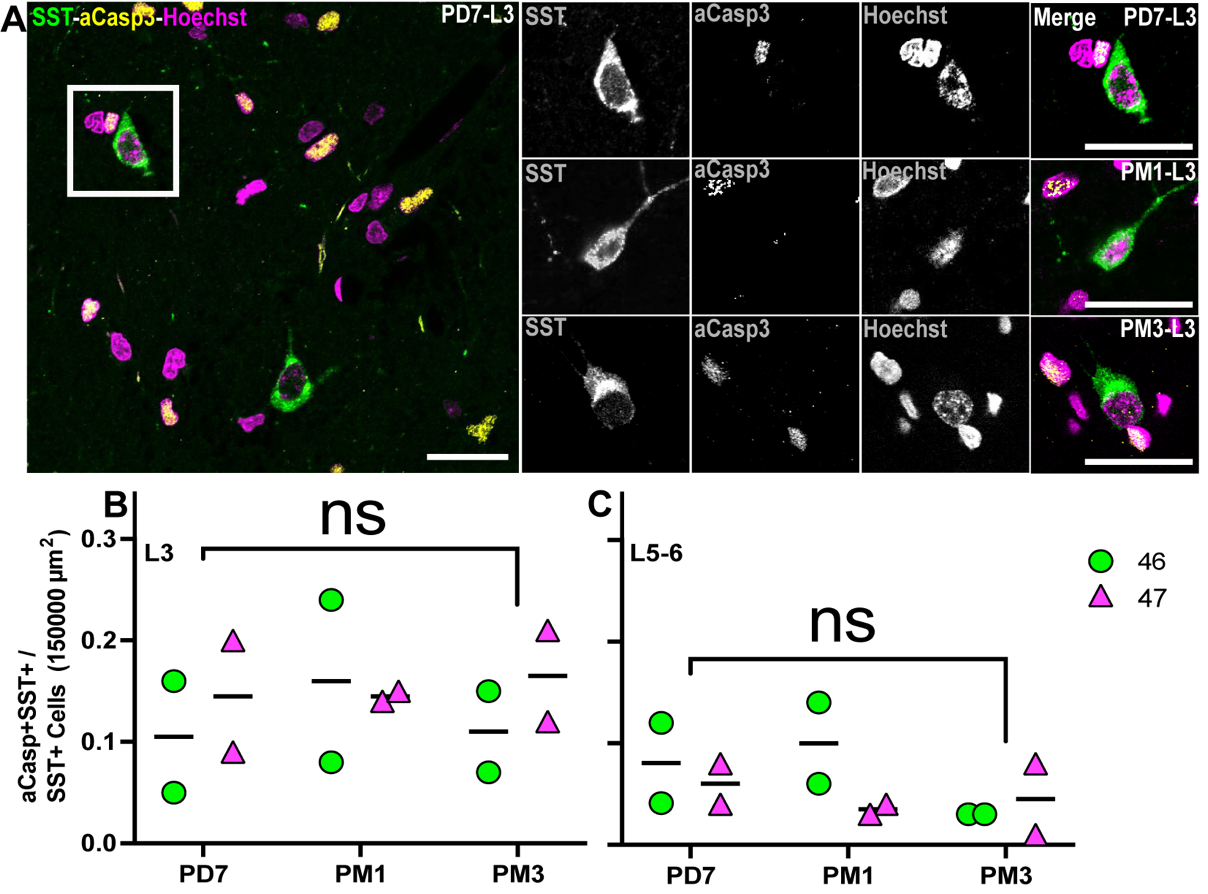

**S1:** **The decrease in the density of SST+ cells did not occur due to apoptosis.**

**A** The images represent triple staining of SST (green), aCasp3 (yellow) and nucleus (magenta) at magnification 60x in L3 of areas 46 at the stages of PD7, PM1 and PM3. **B** and **C** The density of aCasp3+SST+ cells was counted in L3 and 5/6 of areas 46 and 47L at a magnification of 20x; the data analysis did not reveal a significant change in the population of aCasp+ SST+ interneurons in the studied areas suggesting that the postnatal decrease in the fraction of SST+ cells did not happen due to developmental apoptosis. For data analysis, Dunn’s multiple comparisons accompanied the nonparametric Kruskal-Wallis test. The data were presented as median ± interquartile ranges, p<0.05: significant, ns: non-significant (n=2).

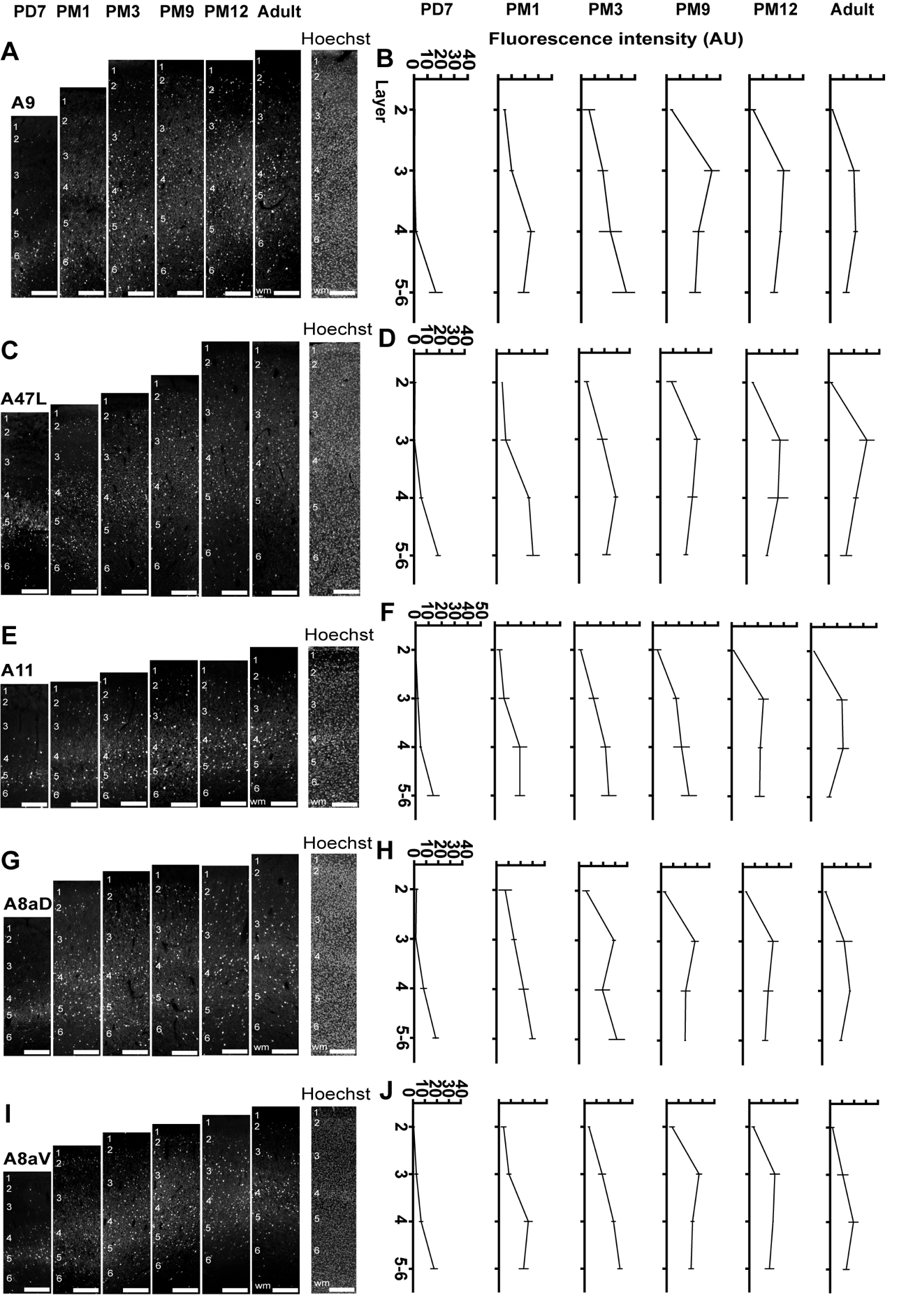

**S2: Consistent modifications of PV laminar distribution across prefrontal areas.**

**A, C, E, G,** and **I** The immunostainings show postnatal developmental modifications of PV (grey) across the laminae in areas 9, 47L, 11, 8aD, and 8aV, respectively. **B, D, F, H,** and **J** The graphs consistently show the quantifications of PV fluorescence intensity in areas 9, 47L, 11, 8aD, and 8aV, respectively. The populations of PV+ interneurons were initially observed in infragranular layers at PD7 and gradually increased towards supragranular L3.

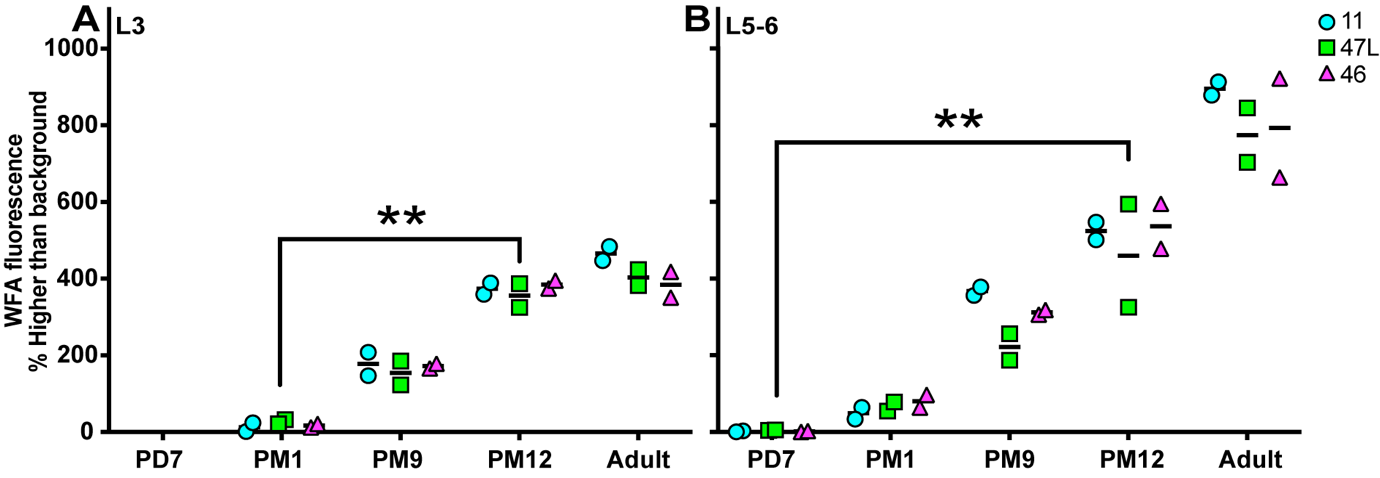

**S3: Consistent increase in PNN fluorescence intensity across prefrontal areas during mid-adolescence.**

**A,** and **B** Graphs represent a gradual postnatal increase in the fluorescence intensity of WFA in the cell membrane of PV+ cells in L3 and 5/6 in areas 11, 47L and 46 at PM12 correlated with mid-adolescence. The amounts of WFA FI were normalized by the autofluorescence intensity of the background. For data analysis, Dunn’s multiple comparisons accompanied the nonparametric Kruskal-Wallis test. The data were presented as median ± interquartile ranges, * p<0.05: significant (n=2).
